# Periodic boundaries in Molecular Dynamics simulations: why do we need salt?

**DOI:** 10.1101/2022.10.18.512672

**Authors:** Wojciech Kopec, Vytautas Gapsys

## Abstract

Molecular dynamics (MD) simulations are usually performed by employing periodic boundary conditions (PBC). While this treatment of simulation system removes the necessity to treat the interactions with an otherwise artificial boundary, PBC also introduces additional constraints that need to be carefully considered for a robust and reliable simulation. Some of the issues pertaining to PBC are well explored and can be remedied by choosing a large enough unit cell, or by applying corrections to the generated trajectories. In current work, we study another artifact which cannot be alleviated by changing the box size. The artifact occurs due to the PBC imposed constraints affecting systems with permanent uncompensated dipoles, which is of particular relevance for lipid membrane simulations. Such dipoles often arise in many biologically-relevant setups, in particular those involving asymmetric lipid bilayers. The artifact manifests itself as an electric field formation in the simulation box which is counteracted by redistribution of mobile charge carriers (ions) and/or ordering of water dipoles. In the absence of ions, the artifact may cause strong water ordering, affecting thermodynamics of the studied system. This observation reveals a conceptually interesting effect of using explicit salt in MD simulations: ions help removing the unwanted periodicity-induced artifact occurring due to uncompensated electric dipoles. Therefore, we recommend adding mobile ions in molecular simulations whenever possible, and call for caution when simulating systems that require low salt concentration (or no salt at all), for example ion channel inactivation promoting conditions. In general, our findings are relevant for molecular simulations of any systems that contain uncompensated dipoles, that might occur more often than previously thought.

## 1 Introduction

Molecular Dynamics (MD) simulations of complex biological and material systems are nowadays routinely used to characterize their behavior on an atomistic scale. Throughout the years of method development numerous best practices have been set ^1–3^ to ensure robust performance of simulations to reliably reproduce physical behaviour of (bio)molecules. The development of these best practices usually involves first an observation of a unexpected and/or unphysical behavior of a given system, due to a spurious combination of system properties and simulation settings, followed by the optimal settings or algorithmic corrections to avoid such artifacts in future simulations. Some examples involve the flying ice cube artifact ^4^, directional flows in the absence of a driving force ^5^ and electrostatics induced artifacts due to non-neutral simulation box with Ewald summation ^6^. One source of potential artifacts is the common practice in MD simulations of using periodic boxes (periodic boundary conditions (PBC)) to approximate bulk behavior of the system of interest. Such a setup alleviates the complications of treating interactions at a boundary of the simulation box. However, this comes at a cost: the periodicity of the system itself imposes a certain degree of artificiality which needs to be considered for generating reliable statistical ensembles.

For example, proper treatment of electrostatics is particularly important due to the long range nature of these interactions. Typically, electrostatics is computed by means of the Ewald summation method (e.g. particle mesh Ewald (PME) ^7^). When a periodic box coupled with PME contains an overall uncompensated electric charge and inhomogeneous dielectric, severe artifacts might arise, for example a stabilization of an ion in the middle of a low dielectric medium (e.g. lipid membrane). See Hub et al. ^6^, for the full discussion.

Recently, we have observed ^8^ a different periodicity induced artifact, which, to our best knowledge, has not yet been explored in detail in the context of MD simulations. We noticed that the periodicity related electrostatic artifacts can occur also in a neutral simulation box, when the box contains an uncompensated electric dipole. Namely, we observed a water ordering effect ^8^ (non-zero values of the average of the cosine of the angle between the water dipole moment and z-axis of the box, ⟨cos(Θ)⟩,A) in the regions far away from the bilayer) (Fig 1 A) in some of our simulations. This effect was introduced by a strong dipole created across a lipid bilayer: the bilayer contained 29 negatively charged POPS lipids in each leaflet, while a polycation carrying 40 positive charges was adsorbed onto one of the leaflets. The overall charge of the box was kept neutral by addition of 18 potassium cations.

**Figure 1:**
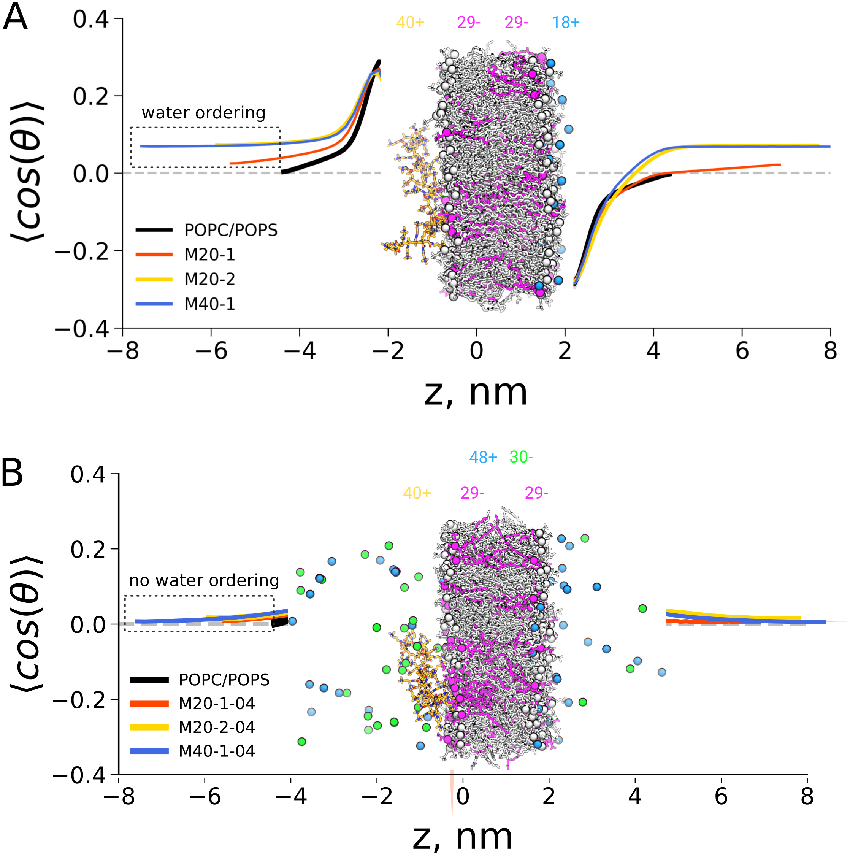
Water ordering in a simulation box with an uncompensated dipole induced by a polycation (orange) binding to the membrane (grey, magenta). Potassium ions are shown as blue spheres. Ionic charges carried by polycation, negatively charged POPS lipids and potassium ions are highligted (A). Individual lines refer to either pure membrane system (black), systems with a shorter polycation of 20 units with either one (M20-1) or two (M20-2) polycation molecules and a system with one polycation with 40 units (M40-1). See our previous work ^8^ for details. This water ordering effect disappears upon the addition of salt (B).

As the polycation was effectively immobilized on one of the leaflets during an entire simulation, the positive charges were separated across the membrane, creating a strong, uncompensated dipole. This way separating +11/-11 charges on the opposite leaflets of the bilayer corresponds to ∼ 3V voltage across PBC in case of M40-1 polycation setup. Were the voltage of this magnitude to drop across the lipid bilayer, it would quickly disrupt the membrane, yet with the periodic conditions applied the high permittivity dielectric, water, reacts by ordering its dipole moments along the direction of the electric field. After observing this artifact, we alleviated it by adding 48 potassium cations and 30 chloride anions (salt), reasoning that mobile charged particles would counteract this strong dipole. Indeed, in all simulations with the added salt, the water ordering disappeared, whereas the polycation remained bound to the membrane. (Fig 1 B).

A schematic way to illustrate this effect is depicted in Fig 2. Here, a lipid bilayer contains leaflets with asymmetric lipid composition: one leaflet is enriched with positively charged lipids, while the other contains negatively charged molecules. If the simulation timescale does not allow for lipid flip-flops between the leaflets, such setup will create permanently separated charges across the membrane. The effect of this charge separation becomes clear when shifting point of view by translating the unit cell along the z-axis (right panel in Fig 2): PBC ensure translational invariance of the system. In this view, it is evident that the simulation box contains fixed charges at its opposite edges. The electric field between the charges is countered by water molecules orienting their dipole moments accordingly. Therefore, this structuring of water is a clear manifestation of a periodicity induced artifact.

**Figure 2:**
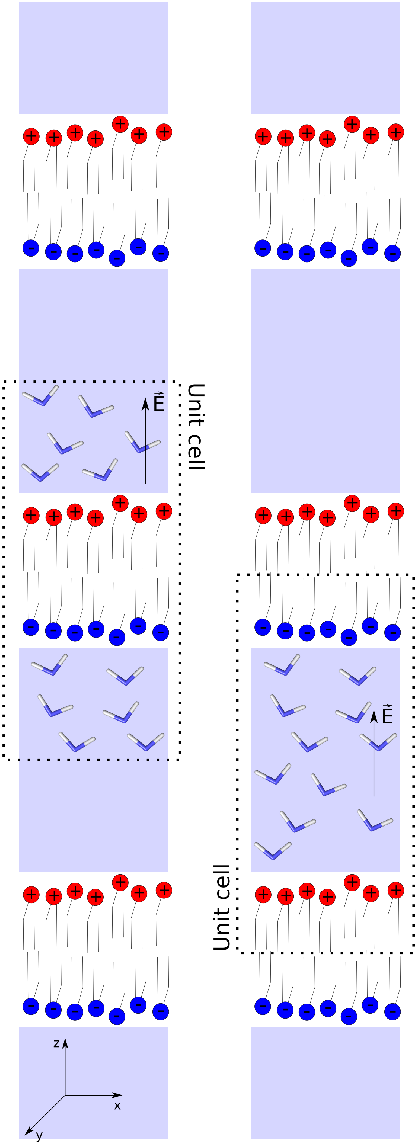
Schematic representation of the periodic environment along z-axis for a single bilayer system. A bilayer consisting of positively (blue) and negatively (red) charged lipids surrounded by water (iceblue), in a periodic box, is shown. The system remains electrically neutral, but the charge separation creates an electric dipole, that is countered by reorientation of water molecules. Translating a unit cell along the z-axis (due to the PBC) illustrates that this situation is equivalent to two charged layers at the edges of the box, with water in between. In this view, it is clear that water dipoles would reorient themselves due to the electric field created by the charged layers.

Another way to explain this effect is by considering that a dipole in the membrane with a fixed orientation would create a difference in potential between the compartments separated by the bilayer. However, the different compartments are connected via PBC, thus making them the same compartment (again, translating the unit cell helps to illustrate this, see Fig 2), thus imposing no potential difference. Yet, what happens to the potential difference which was created by the fixed dipole of the asymmetric bilayer? In this case, the PBC can be viewed as a constraint which creates another dipole to counteract the effect of the bilayer. In fact, this explanation is more general than the example with separated charges in Fig 2), because not only asymmetric charge distribution, but also asymmetry in lipids of different chemistry between the leaflets can lead to the potential difference across the membrane ^9^.

The emergence of such an electric field due to PBC has been described in Quantum Mechanical Density Functional Theory (DFT) calculations using plane-wave methods. ^10,11^ To correct for this artifact in DFT, a solution was suggested of explicitly adding a counteracting dipole field. In the field of MD simulations, the effect of simulation cell dipole has been noted in the context of electrostatic interaction calculation by means of Ewald summation. ^12,13^ It has been observed that the true electrostatic energy of a periodic system differs from that calculated by means of the Ewald summation by a factor proportional to the squared dipole moment of the system in the unit cell. This extra term vanishes when the macroscopic system (infinite periodic cells) are surrounded by a conductor, i.e. tin-foil boundary. However, neither using the tin-foil boundary, nor a different insulator resolves the artifact in MD simulations, as we will show in the current work. A possible pragmatic solution to the problem is using mobile charge carriers, i.e. ions, which allow to alleviate the artifact by redistributing the charges to counteract the electric field.

In the current work, we investigate and quantify the effects of uncompensated electric dipoles in MD simulations of biological systems. By considering several test systems, we show how the uncompensated dipoles trigger either a counter-dipole formation through ordering of water molecules in the bulk. If two of such dipoles are inserted in the same simulation box in a way that they directly compensate each other (e.g. in a double membrane setup), a formation of explicit electric potential is observed (instead of water ordering), which provides an intuitive way of quantifying the effect. In all cases, these artifacts are removed by an addition of salt (mobile ions), in high enough concentrations, to bulk water. As the asymmetry of biological membranes, in terms of lipid composition and the shapes of membrane proteins, is widespread ^14^, and the processes happening at low salt concentrations (e.g. ion channel inactivation ^15^) are biologically relevant, we believe that our contribution will help avoiding artifacts in MD simulations of such systems. More generally, our findings apply to all simulations in periodic boxes with uncompensated electric dipoles, and thus can be useful also for e.g. simulations of inorganic surfaces or other materials with exotic shapes.

## 2 Methods

All MD simulations reported in this work are listed in the 1) and have been performed with the GROMACS simulation package, versions 2018-2020. The POPE/POPC, POPC/DOTAP/POPS and POPC (with and without vacuum slabs) membranes were generated using the CHARMM-GUI web interface ^16^. Systems containing the NaK2K channel were prepared based on NaK2K simulations from our recent paper ^17^ and the desired protonation state was set manually. For the free energy setup (anthracene with the POPC/DOTAP/POPS membrane) we used the previously generated structures and topologies from our previous work ^18^.

All simulations were run using the CHARMM36m force field for proteins and lipids ^19^, with the TIP3P water model ^20^ and default CHARMM ion parameters ^21^. CGenFF parameterization was used for anthracene ^22,23^. All hydrogen-containing bonds were constrained using LINCS ^24^, allowing for the integration timestep of 2fs. The Particle Mesh Ewald (PME) was used for for electrostatic interactions ^7,25^, with the 1.2 nm real-space cutoff. For the simulations treating electrostatics with PME and relative conductivity of the surrounding boundary of ∈ = 80, the real-space cutoff was set to 1.3 nm. The Nose-Hoover thermostat ^26,27^ and Parrinello-Rahman barostat were used ^28^. Van der Waals interactions were force-switched off from 0.8 to 1.2 nm. Free energy calculations were performed by means of free energy perturbation (FEP). The charges on anthracene were set to zero and the van der Waals interactions with the rest of the system were alchemically switched off using 11 equidistantly spaced *λ* windows.

The analysis was conducted using Gromacs tools: gmx h2order (water orientation), gmx potential (membrane potential), gmx density (ion densities). Free energy was calculated using Bennet’s acceptance ratio estimator ^29^ using the Alchemical Analysis tool. ^30^ Each simulation copy for the same system (see Table 1) has been treated as an independent measurement, to calculate averages and their error bars (standard error of the mean). Visualisations were done with VMD ^31^, numerical analysis and plotting with numpy ^32^ and matplotlib ^33^, respectively.

**Table 1:** Summary of the simulations performed in this work.

| System | Notes | Number of simulations<br>& simulation time |
| --- | --- | --- |
| POPE/POPC<br>single membrane | 1 system without salt | 3 x 250 ns |
| POPE/POPC<br>double membrane | 2 systems: without salt and with -0.35/+0.35 charges | each system 3 x 250 ns |
| POPC/POPC<br>double membrane | 1 system without salt | 3 x 250 ns |
| POPC/DOTAP/POPS<br>single membranes | 10 systems without salt with<br>PME tin-foil, cut-off and<br>10 systems with 100 mM KCl (PME tin-foil):<br>0, 1, 2, 3, 4, 5, 6, 8, 10, 12 charged lipids per leaflet | each system 3 x 250 ns |
| POPC/DOTAP/POPS<br>single membranes | 10 systems without salt with PME $\epsilon = 80$:<br>0, 1, 2, 3, 4, 5, 6, 8, 10, 12 charged lipids per leaflet | each system 3 x 80 ns |
| POPC/DOTAP/POPS<br>double membranes | 3 systems (all without salt):<br>0, 1, 2 charged lipids per leaflet | each system 3 x 250 ns |
| Anthracene +<br>POPC/DOTAP/POPS<br>single membrane | FEP calculations<br>10 systems without salt:<br>0, 1, 2, 3, 4, 5, 6, 8, 10, 12 charged lipids per leaflet | each system 5 x 10 ns<br>11 $\lambda$ windows |
| NaK2K channels +<br>POPC single membrane | 12 systems:<br>without salt, with 5-50 mM KCl, with 150 mM KCl | each system 3 x 1000 ns |
| POPC/POPC<br>single membrane<br>separated charges | 2 systems with separated 10 Na<sup>+</sup> and 10 Cl<sup>-</sup> ions:<br>setup with vacuum slab and<br>setup with ions restricted to the xy-plane | each system 5 x 50 ns |

## 3 Results

### Membrane asymmetry induces electric dipoles

We started our investigation with a simple asymmetric POPE/POPC membrane, that is with POPC lipids in one leaflet and POPE lipids in the second one (Fig 3 A), inspired by the previous work where such a lipid asymmetry created a potential difference across the simulation box ^9^. This system allows us to quantify the effect of an overall dipole with a fixed orientation on the potential difference across the box and water ordering in a biologically relevant setup, without the need of charge separation.

**Figure 3:**
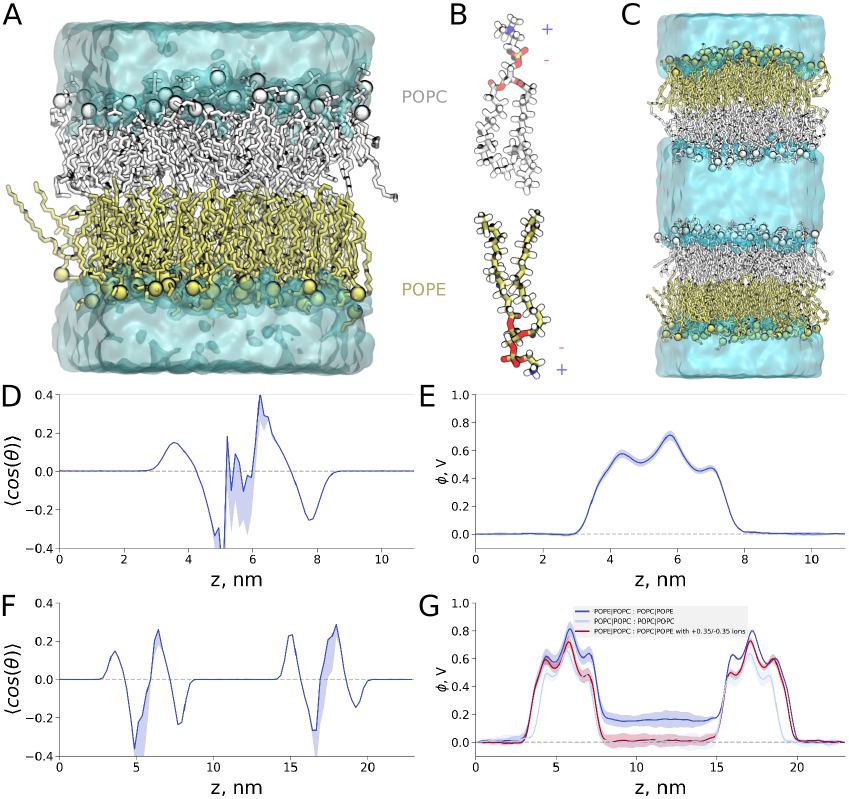
Artifacts in asymmetric POPE/POPC membranes in the absence of salt. (A) Visualisation of the single membrane system, made of POPC (grey) and POPE (tan) lipids. Water is shown as a cyan surface. (B) A zoom-in on the individual lipid molecules, showing the dipole formed their headgroups. (C) The double membrane systems with two oppositely oriented asymmetric POPE/POPC membranes. No water ordering in the bulk is observed in either (D) single membrane or (F) double membrane setup. (E) The potential difference between the water bulk regions in the single membrane setup is 0 V. (G) The double membrane setup reveals that POPE/POPC bilayer asymmetry creates potential difference of 0.2 V between the compartments (dark blue curve). This potential corresponds to the separation of +0.35/-0.35 charges: explicitly adding counter ions with such charges removes the potential difference (dark red curve). For a symmetric POPC/POPC membrane the potential difference between the compartments is 0 V (light blue curve).

Both POPE and POPC lipids are electrically neutral, but form electric dipoles due to their zwitterionic character (Fig 3 B). The shape of the lipid bilayer and a large energetic barrier for the exchange of lipids between leaflets (lipid flip-flop) limits the mobility of POPC and POPE molecules to their respective leaflets, creating an overall electric dipole, which remains uncompensated throughout the microsecond timescale of our simulations. Since periodic simulation system constrains potential difference across the box to 0 V, we also constructed a double membrane setup, by oppositely orienting POPE/POPC bilayers and placing them in the same box (Fig 3 C). In this setting, we are able to quantify the effect of asymmetric membrane dipoles by monitoring the electric potential difference between the water compartments separated by the bilayers. Interestingly, despite the presence of an overall uncompensated dipole, no water ordering in the bulk was detectable in the single membrane setup (Fig 3 D). As expected, the potential difference across the box was 0 V (Fig 3 E). The surprising lack of the water ordering effect is in contrast to the polycation-bilayer system (Fig 1), therefore, to understand the cause of this discrepancy, we investigated the double bilayer setup of the POPE/POPC membrane. Expectantly, no water ordering is observed for this setup as well (Fig 3 F), as the dipole of one bilayer compensates the dipole of the other one. Yet, for the double membrane system, we are able to quantify the potential difference induced by the leaflet asymmetry (Fig 3 G). The membrane potential difference reaches a relatively low value of 0.2 V, which can be counteracted by adding a single pair of oppositely charged particles (ions) with +0.35/-0.35 charges to the separated water compartments (Fig 3 G, dark red curve). This observation provides a simple explanation of the absence of observable water ordering in the asymmetric POPE/POPC bilayer setup, as it is a consequence of a much smaller overall dipole than in the polycation-membrane system, where the charge difference between the leaflets was -11/+11. This observation naturally prompts the question: at what level of the membrane asymmetry (dipole strength) is the water ordering artifact detectable?

### Systematic investigation of the artifacts

To control the electric dipole of a lipid membrane we constructed a POPC/POPC bilayer with a varying number of negatively charged POPS lipids in one leaflet and positively charged DOTAP lipids in another leaflet ((Fig 4 A, B).

**Figure 4:**
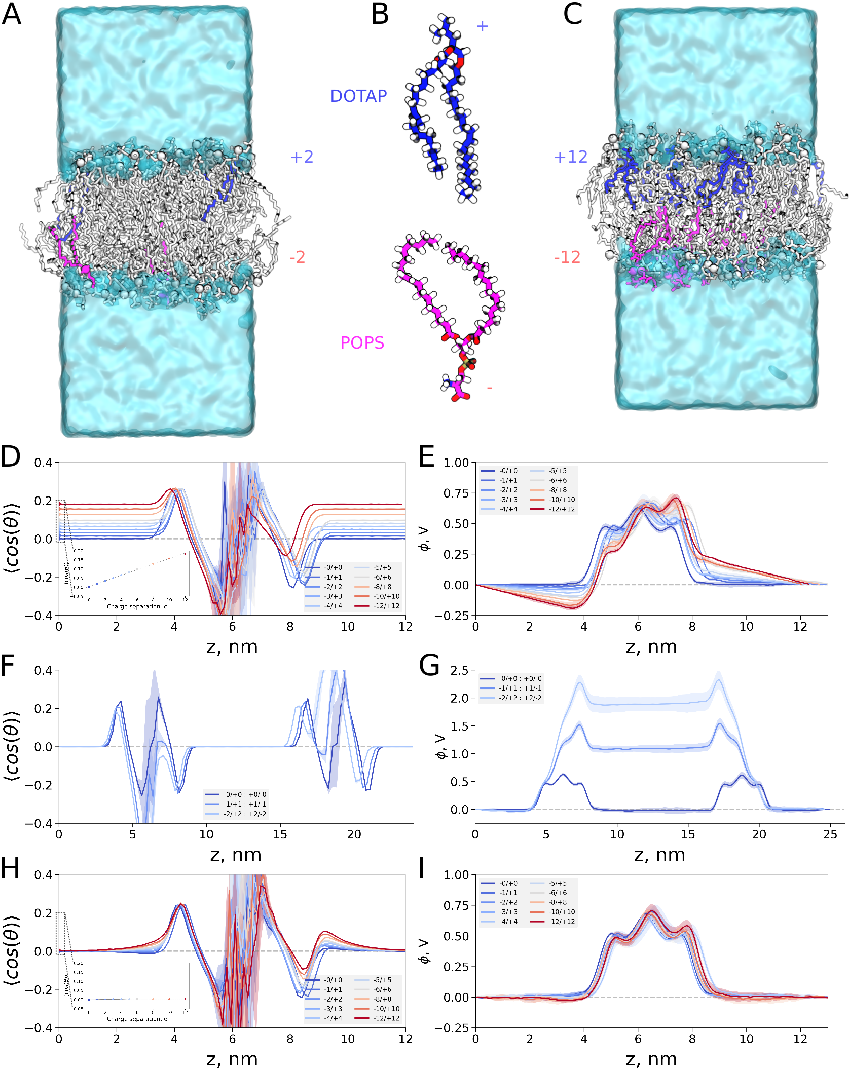
Artifacts in asymmetric POPC/DOTAP/POPS membranes with separated charges in the absence of salt. (A) Visualisation of the single membrane system, made of POPC (grey), two cationic DOTAP (blue) and two anionic POPS (magenta) lipids. Water is shown as a cyan surface. (B) A zoom-in on the individual DOTAP and POPS molecules. (C) Same as (A), but with 12 DOTAP and 12 POPS molecules. (D) Averaged cosine of an angle (⟨cos(Θ)⟩) between the water dipole moment and normal to the membrane in the single bilayer setup. The leaflets contained varying number of negatively (POPS) and positively (DOTAP) charged lipids. The inset shows the ⟨cos(Θ)⟩ values in the bulk as a function of the number of separated charges. (F) This water ordering is absent in the double membrane setup. (E) The potential difference between the water bulk regions in the single membrane setup is 0 V. (G) The potential difference across the simulation box in the double membrane setup. The addition of mobile ions (salt) removes water ordering (H) at all charge separation (H, inset) and makes the membrane potential symmetric (I) in the single membrane setup.

By systematically inserting more and more (1-12) POPS and DOTAP molecules into the opposing leaflets (Fig 4 C), we increased charge separation across the membrane, thus inducing progressively larger electric dipoles. Here, we again used a single membrane setup to monitor the magnitude of water ordering artifact, while the double bilayer setup allowed us to relate the varying charge separation to the changes in the membrane potential.

A small, yet significant bulk water ordering is visible already with the separation of 2 charges (−1/+1) (Fig 4 D). With the larger charge separations the water ordering increases linearly (inset in Fig 4 D), which is also clearly visible when considering the average dipole moment of bulk water molecules Fig 5). Similarly to the previous system, in the single bilayer setup, however, we again cannot quantify the membrane potential these dipoles would create, as the periodic boundary conditions constrain the potential difference across the box to 0 V Fig 4 E).

**Figure 5:**
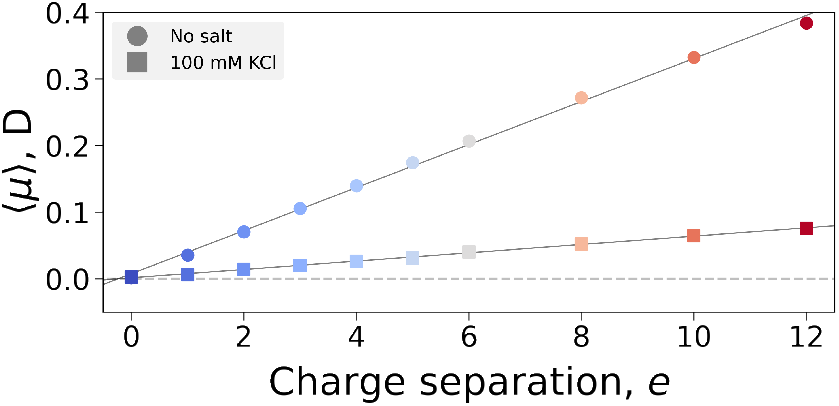
Average bulk water dipole moment per water molecule for systems containing lipid bilayer with asymmetric distribution of DOTAP/POPS lipids in the leaflets.

Such a water ordering effect has an immediate effect on thermodynamics of solvated molecules. As an illustration, we have calculated hydration free energy of an organic molecule anthracene in each of the systems analyzed in Fig 4. The calculation was carried out by placing anthracene in the bulk water away from the membrane. One atom of anthracene was position restrained to ensure that the molecule remained away from the membrane: the minimal distance to any bilayer atom was kept larger than the non-bonded interaction cut-offs. In Fig 6 we show one component to the hydration Δ*G*, namely contribution from coupling the van der Waals interactions of anthracene to water. Analyzing only the Δ*G*_*vdw*_ component makes it possible to assess the effect of water ordering only, as the full hydration free energy contains also the electrostatic component Δ*G*_*Q*_, which is inevitably affected by altering the charges on the membrane. Fig 6 shows a clear and significant influence of the artifact on thermodynamics of anthracene solvation.

**Figure 6:**
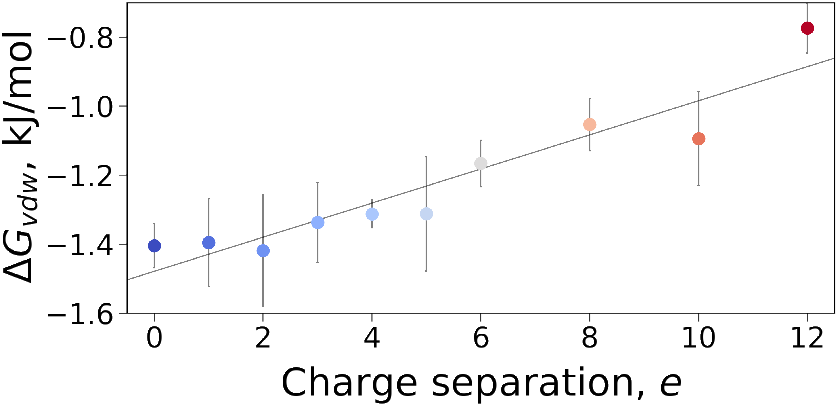
Van der Waals interaction component Δ*G*_*vdw*_ of the anthracene hydration free energy in bulk water of systems containing lipid bilayer with asymmetric distribution of DOTAP/POPS lipids in the leaflets. Δ*G*_*vdw*_ depends on charge separation between the membrane’s leaflets.

The artifact introduced by separating charges between the bilayer leaflets remains irrespective of the electrostatic interaction treatment. In particular, we observed the same water ordering effect when changing the relative permittivity of the conductor surrounding infinite periodic system (Fig 7). In Fig 4 we performed simulations where electrostatic interactions were treated with Particle Mesh Ewald (PME) using a conductor to surround the infinite periodic system, i.e. tin-foil boundary conditions, ∈ = ∞. Fig 7A Demonstrates that changing the relative permittivity of the boundary (∈ =80) has no effect on the artifact. We have also probed whether this PBC effect can be in general attributed to the PME artifacts. For that we disabled PME and relied on the plain cutoff electrostatic calculations: while such a simulation would not yield an overall rigorous thermodynamic ensemble, it is meaningful for testing whether the artifact remains. The average water orientations for the cutoff simulations (Fig 7B), as expected, show some difference from the PME results, yet significant water ordering effect remains.

**Figure 7:**
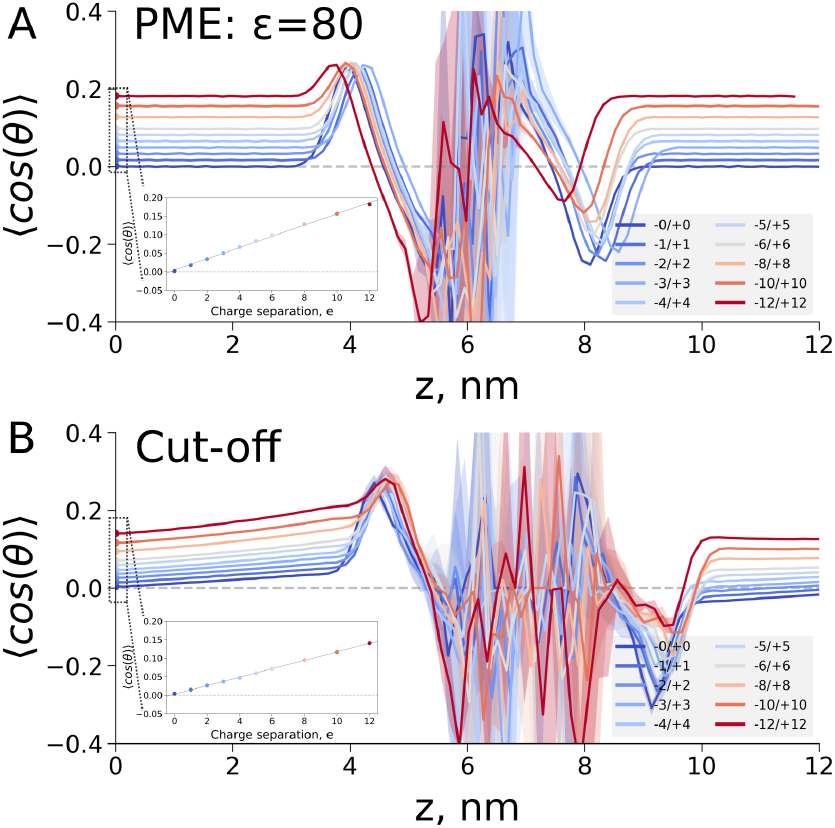
Averaged cosine of an angle between the water dipole moment and normal to the membrane, ⟨cos(Θ)⟩, in the single bilayer setup with different numbers of DOTAP/POPS lipids in the leaflets. (A) Electrostatics were treated with PME and tin-foil boundary conditions. (B) Plain cutoff was used for electrostatics.

We have also used the double bilayer setup, where we do not observe any artifact of bulk water ordering (Fig 4 F), but, in turn, we clearly see that already separation of 8 charges (−2/+2 : +2/-2 in the double bilayer, which corresponds to the -4/+4 separation for the single bilayer setup) introduces a huge membrane potential of 2 V. (Fig 4 G). This voltage is high enough to induce occasional membrane poration within 250 ns MD simulations, in line with previous reports ^34^. Therefore, larger charge separations in the double bilayer setting are not feasible, as the created large potential difference quickly disrupts the membrane.

Our systematic study of the charge separation between the lipid bilayer leaflets highlights the magnitude of an artifact that may potentially be unnoticed in simulations with a permanent uncompensated dipole of the single lipid membrane in a periodic box. Induced high voltages, otherwise capable of disrupting the membrane, will be compensated by orienting dipoles in the rest of the system to fulfill the periodicity imposed constraint of 0 V between the ends of the periodic box. These large artifacts can be alleviated (Fig 4 H,I, Fig 5) by supplying the system with mobile charges (i.e. salt) that could dynamically redistribute to compensate the membrane dipole (Fig 8).

**Figure 8:**
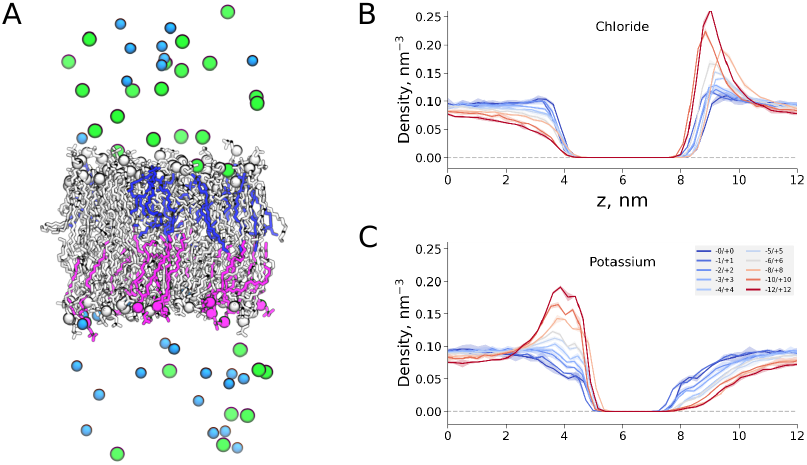
Density of ions (chloride and potassium in simulations of a single lipid bilayer with separated charges. (A) Visualisation of the system, POPC lipids are shown in grey, POPS in magenta and DOTAP in blue. Chloride ions are shown as green spheres and potassium ions are light blue spheres. (B) and (C) density of chloride and potassium ions, respectively, across the simulation box.

So far we have mainly focused on the asymmetries imposed by the membrane composition or large molecule adsorption at the membrane surface. Next, we noticed that the integral membrane proteins also introduces asymmetries in biological membranes, potentially leading to artifacts in MD simulations of these systems.

### Membrane asymmetry due to an embedded protein

A great majority of integral membrane proteins (i.e. embedded in the membrane) are asymmetric across the membrane, and often decorated with positively (arginine (arg), lysine (lys)) and negtively (glutamate (glu), aspartate (asp)) charged residues, and thus capable of inducing a charge asymmetry across the membrane. Here, we used a tetrameric potassium channel NaK2K (Fig 9), that has in total 12 asp residues (all on the extracellular side), 8 glu residues (4 on the extracellular side, 4 on the intracellular side), 4 arg residues (all on the extracellular side) and 8 lys residues (4 on the extracellular side, 4 on the intracellular side).

**Figure 9:**
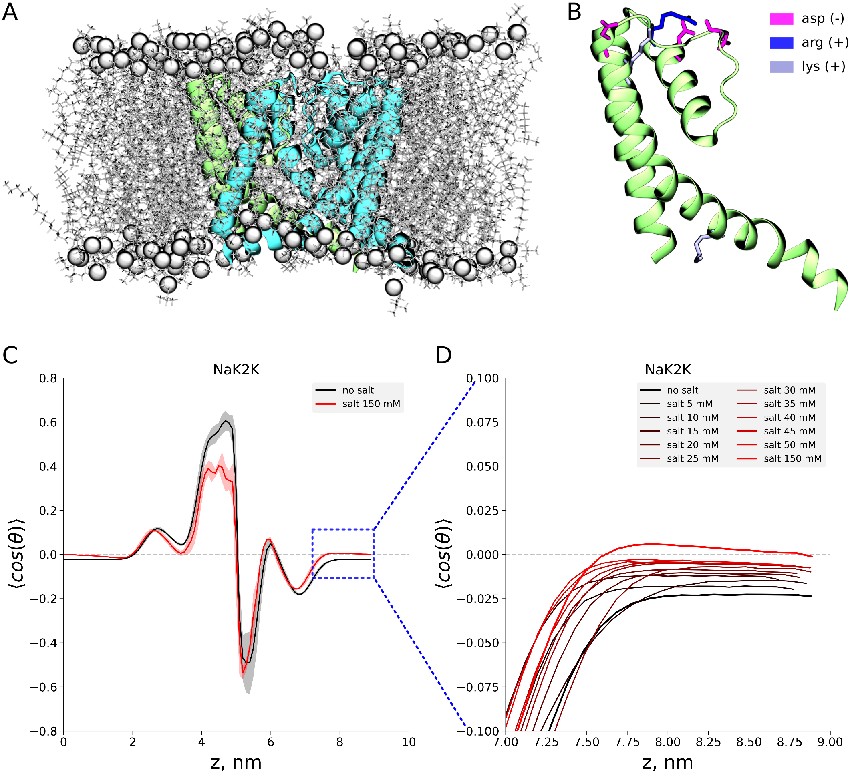
Water ordering caused by the membrane embedded protein. (A) A tetrameric potassium channel NaK2K (cyan) embedded in a POPC membrane (grey). Charged residues in NaK2K are shown for a single monomeric unit (B), highlighting their asymmetric distribution across the membrane. Such a charge distribution across the membrane induces water ordering (C), which can be removed by the addition of salt (D), but only in high enough concentration.

If these residues were assigned to their standard protonation states at pH 7 (i.e. all asp and glu residues would be negatively charged, and all arg and lys residues would be positively charged), the NaK2K channel would have a 8 overall charge on the extracellular side, and 0 on the intracellular side, resulting in the total charge of -8. Such a situation would create a large charge asymmetry across the bilayer (precisely what we want to study), but at the same time would lead to an electrically nonneutral box in a simulation in the absence of salt. Therefore, we decided to protonate all glu residues in our system (Fig 9 B), which makes the system neutral and, at the same time, maintains a strong dipole across the membrane (−4 overall charge on the extracellular side and +4 overall charge on the intracellular side). This way, we study a system similar to one induced by polycation binding 1, however now created solely by a distribution of charged residues in a membrane embedded protein, thus relevant for many biological systems. When this system is simulated in the absence of salt, the expected water ordering behavior is observed, which disappears upon addition of 150 mM salt (KCl) (Fig 9 C). However, simulations at lower salt concentrations (Fig 9 D) revealed that some water ordering persists even at 50 mM salt concentration (corresponding to 10 K+ and 10 Clions in the simulated system), thus one needs to be careful when simulating membrane proteins at relatively low salt concentration.

## 4 Discussion

The artifact that we investigated in the current may affect many molecular dynamics simulations setups. Namely, it is created by permanent dipoles pertaining in the system throughout the simulation time. While in the work we show examples of lipid membrane to illustrate the artifact, other systems may also be affected: for example, permanent dipole moments in bio-molecular simulations may be imposed by restraining the molecules or removing angular center-of-mass motion ^35^. Similarly, for the molecules with slow rotational diffusion, the simulation time may not be sufficient to exhaustively sample molecular orientations, thus retaining the direction of the dipole unchanged throughout the simulation. This may be of importance for simulating large bio-molecules at relatively short time scales. ^36,37^ Furthermore, the described periodicity effects might be of concern for simulations of the solid state, surfaces and interfaces in material sciences ^38–42^

The manifestation of the artifact can be alleviated by explicit addition of mobile charge carriers, ions, to the simulation box. Mobile ions redistribute to counteract the electric field formed due to the periodic boundary conditions. This way, water dipoles or other molecules in the bulk are not affected by the artifact. The number of ions needed varies, e.g. for the asymmetric POPE/POPC bilayer addition of two ions with the +0.35/-0.35 charges was sufficient to remove the otherwise created potential difference (Fig 3). However, for the NaK2K channel example, where +4/-4 charges were separated across the membrane (Fig 9) it was required to reach physiological salt concentrations of 150 mM to remove the water ordering effect.

Naturally, one needs to consider the possible effects of this artifact when simulating in low salt concentrations or without salt at all. Even though explicit addition of ions to reach physiological salt concentration is frequently adopted practice nowadays, certain studies may require low salt concentrations (or no salt addition at all) ^15,43–48^. On top of that, one needs to consider what low salt concentration might mean at the small scales used in MD simulations. In experiments, even at low salt concentrations, a certain number of ions is expected to be close to the membrane surface (or a protein), as well as in the bulk water. In small boxes used in MD simulations however, the same nominal low salt concentration might mean, in practical terms, only a handful of ions being present, which might neither be enough to fully remove the water ordering artifact, nor be representative of the experimental situation.

Overall, this finding reveals an unexpected benefit of using explicit counter ions in the molecular dynamics simulations. Addition of physiological salt concentration is generally advised by the best practice guides in MD simulation setup. ^49–51^ An intuitive explanation for this requirement relies on the need to accurately represent specific interactions that might occur in the experimental setup. However, as we have demonstrated, from a conceptual point of MD simulations, there is a need for mobile charges in a periodic system. By altering ion distributions, mobile counter ions compensate for potential artifacts that might occur due to dipoles with a fixed orientation in the simulation box.

While the described artifact itself is of electrostatic nature, it is not dependent on the particular method for treating electrostatic interactions (Fig 7) and manifests due to the constraints imposed by the periodic boundary conditions. In fact, this effect of electric field created by PBC was played an important role for methods investigated in multiple publications previously, only it was not directly identified and named as an artifact. ^9,52–54^ In numerous studies a transmembrane potential was simulated by creating a system with a lipid bilayer allowing for a vacuum slab to split the bulk water layer along z axis (Fig 10A). It is an intuitive way to separate the charges on the opposite sides of the bilayer, thus generating transmembrane potential. For example, in Fig 10A we have prepared a system with 10 Na^+^ on one side of the membrane and 10 Cl^−^ ions on the other side. In turn, the created voltage is so high that once the simulation starts, the membrane quickly porates (Fig 10B).

**Figure 10:**
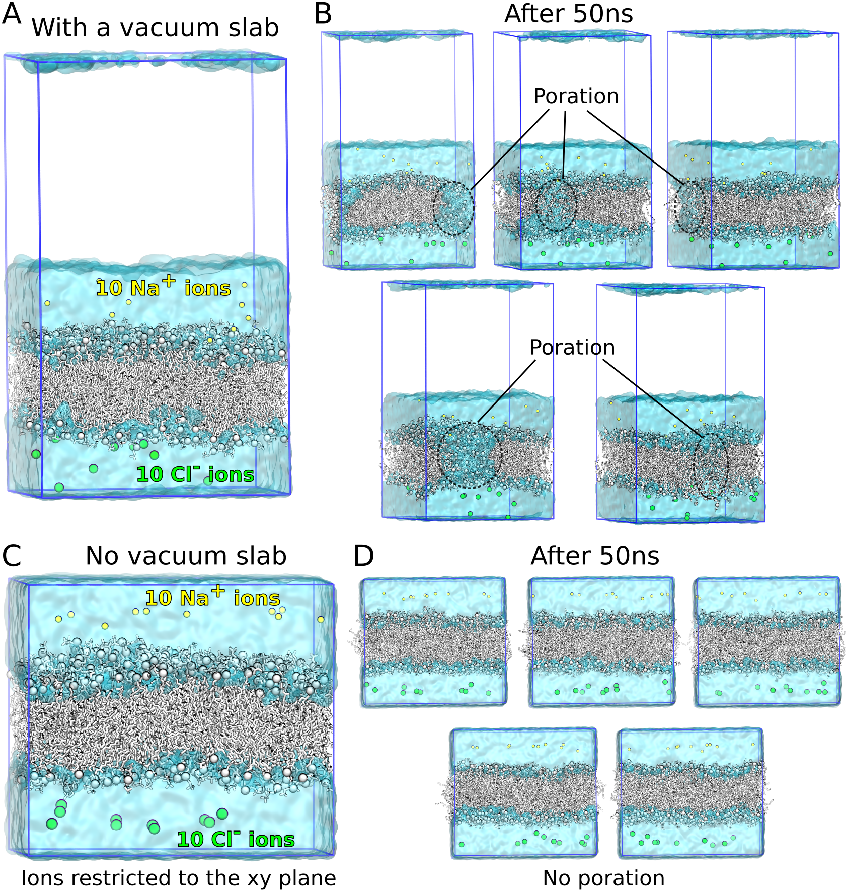
Phospholipid bilayer (POPC/POPC) setup for MD simulations with separated charges. (A) Vacuum slab splits the bulk water layer along z-axis. 10 sodium and 10 chloride ions are separated on the opposite sides of the bilayer. (B) Snapshots from 5 independent simulation replicas after 50 ns with the membrane poration regions marked. (C) Phospholipid bilayer without vacuum slab to interrupt the bulk water layer. Sodium and potassium ions are restricted to the xy-plane preventing them from moving along z-axis. (D) No membrane poration occurs in any of the 5 independent replicas.

Charge separation, however, is not the only effect that the vacuum slab provides. It also creates a layer of low permittivity as an insulator to the electric field across PBC. To demonstrate this we have also prepared a simulation system separating the charges on the opposite sides of the bilayer by preventing their motions along z-axis instead of inserting a vacuum slab ((Fig 10C). In this scenario, even though we have ensured the same charge separation as in the setup with the vacuum slab, no membrane poration occurs (Fig 10D). The reason for that is possibility for the electric field across PBC to be formed and the high permittivity dielectric (water) counteracts the field by dipole orientation. This also explains why other studies that used similar ion restriction methods, could have created a potential difference only due to the concentration gradient, but not due to the electrostatic potential. ^55,56^

All in all, the external electric field across PBC might have been present in many MD based investigations involving asymmetric lipid bilayers. However, the artifact is mitigated by explicit addition of mobile ions. In cases where no additional salt concentration can be used due to a particular study design, it would be important to ensure whether the artifact could influence the final conclusions of such studies. For that, one might perform test simulations with salt added and assess whether the results are affected beyond the expected explicit interactions of the solute with the included counter ions.

## 5 Conclusion

The best practice of MD simulation recommends neutralizing the system to avoid the artifacts pertaining to the treatment of long range electrostatic interactions. In addition to this, we now add the recommendation of explicit addition of counter ions to avoid the periodicity induced artifacts due to otherwise explicitly uncompensated dipole moments in the simulation box.

